# Human embryonic stem cell-derived cardiomyocytes express SARS-CoV-2 host entry proteins: screen to identify inhibitors of infection

**DOI:** 10.1101/2021.01.22.427737

**Authors:** Thomas L. Williams, Maria T. Colzani, Robyn G.C. Macrae, Emma L. Robinson, Stuart Bloor, Edward J. D. Greenwood, Jun Ru Zhan, Gregory Strachan, Rhoda E. Kuc, Duuamene Nyimanu, Janet J. Maguire, Paul J. Lehner, Sanjay Sinha, Anthony P. Davenport

**Author notes:** Joint first authors, contributed equally (Thomas L. Williams, Maria T. Colzani). Joint senior authors (Sanjay Sinha, Anthony P. Davenport). Correspondence should be addressed to: S.S. and A.P.D. This research was funded in whole, or in part by: Wellcome Trust (WT107715/Z/15/Z, A.P.D., J.J.M.); Addenbrooke’s Charitable Trust (P.J.L., A.P.D.); Wellcome Trust Programme in Metabolic and Cardiovascular Disease (203814/Z/16/A, T.L.W., D.N.), Wellcome Trust Major Award (208363/Z/17/Z) for Imaging Core (G.S.); Wellcome Trust Principal Research Fellowship (210688/Z/18/Z, P.J.L.), UKRI/NIHR through the UK Coronavirus Immunology Consortium (P.J.L.); British Heart Foundation (FS/17/61/33473 APD, S.S., R.G.C.M; TAF 03, APD, JJM); British Heart Foundation Senior Fellowship (FS/18/46/33663, S.S.), British Heart Foundation Centre for Regenerative Medicine (RM/17/2/33380, S.S., M.C.); Cambridge Biomedical Research Centre Biomedical Resources Grant (University of Cambridge, Cardiovascular Theme, RG64226) and a core support grant from the Wellcome Trust and MRC to the Wellcome–MRC Cambridge Stem Cell Institute. For the purpose of open access, the author has applied a CC BY public copyright licence to any Author Accepted Manuscript version arising from this submission.’.

## Abstract

Patients with cardiovascular comorbidities are more susceptible to severe infection with SARS-CoV-2, known to directly cause pathological damage to cardiovascular tissue. We outline a screening platform using human embryonic stem cell-derived cardiomyocytes, confirmed to express the protein machinery critical for SARS-CoV-2 infection, and a pseudotyped virus system. The method has allowed us to identify benztropine and DX600 as novel inhibitors of SARS-CoV-2 infection.

---

The case fatality rate in patients infected with severe acute respiratory syndrome coronavirus 2 (SARS-CoV-2), the emergent cause of the COVID-19 pandemic, rises from 2.3% to 10.5% in individuals with cardiovascular comorbidities^1-3^. Transmission electron microscopy has been used to confirm the presence of SARS-CoV-2 viral particles in human cardiomyocytes infected *in vitro* with the virus, or in autopsy samples from patients positive for SARS-CoV-2. Further, SARS-CoV-2 infection in induced pluripotent stem cell (iPSC)-derived cardiomyocytes induces morphological and cytotoxic effects, and sarcomere disruption in autopsy samples is observed, suggesting SARS-CoV-2 directly damages cardiac tissue^4-6^.

Entry of SARS-CoV-2 into host cells is dependent on the high affinity binding of primed viral spike (S) protein to cell surface angiotensin-converting enzyme 2 (ACE2)^7^. Priming of S protein, through proteolytic cleavage of S1/S2 and S2’ sites is mediated by transmembrane protease, serine 2 (TMPRSS2), also present at the cell surface^8,9^. Further host protein components are implicated in SARS-CoV-2 infection, including the endosomal proteases furin and cathepsins, involved in S1/S2 cleavage and endosomal processing respectively^10,11^. B^0^AT1 (*SLC6A19*) is a neutral amino acid transporter whose surface expression is critically regulated by ACE2. Interestingly, an ACE2-B^0^AT1 heterodimer complex was shown using cryogenic electron microscopy to be able to bind two SARS-CoV-2 S proteins simultaneously^12^. A number of comprehensive reviews outlining potential drug targets for SARS-CoV-2 therapy have been published^13,14^.

We previously demonstrated expression of genes for the proteins listed above in human cardiomyocytes, and have shown they are significantly upregulated in aged patients, providing a rationale for increased susceptibility to SARS-CoV-2 infection in the elderly population^15^. Our aim was to determine whether a human embryonic stem cell-derived cardiomyocyte (hESC-CM) model expresses the same repertoire of recognition, processing, and ancillary genes (and corresponding proteins) as observed in adult cardiomyocytes, and demonstrate viral entry of SARS-CoV-2 in this model. Human stem cell lines used to generate cardiomyocytes are widely acknowledged for their usefulness in cardiovascular medical research, and will be critical in understanding the recognised pathology of SARS-CoV-2 in this tissue type, alongside providing a suitable model for drug screening^16,17^. As a secondary aim, we looked to validate infection of hESC-CMs using a quantitative automated system in conjunction with a pseudotyped HIV-1 based lentivirus decorated with the SARS-CoV-2 spike protein, and screen for novel therapeutic agents that inhibit infection.

We confirm the presence of host cell proteins associated with SARS-CoV-2 infection using immunocytochemistry in the hESC-CM line, and using immunohistochemistry in human left ventricle tissue sections (Fig. 1a-g). This is critical as the expression of receptor recognition and viral entry proteins may vary with cell type. Quantification demonstrated positive immunolabelling above background and control hESC-CMs, with >95% of the population showing immunolabelling for ACE2, TMPRSS2, cathepsin L, and furin (Fig. 1h). B^0^AT1 immunoreactive signal was observed in 27.6% of the cell population, while cathepsin B immunoreactive signal was not observed (0.05% of cells) (Fig. 1h). Results were recapitulated in human left ventricle sections, with immunolabelling detected for all proteins, except cathepsin B which displayed little to no immunoreactivity. Expression of the corresponding genes for the proteins listed above was also demonstrated in hESC-CMs and human left ventricle tissue, except *CTSB* in ventricular tissue (Fig. 1i).

**Fig 1.**
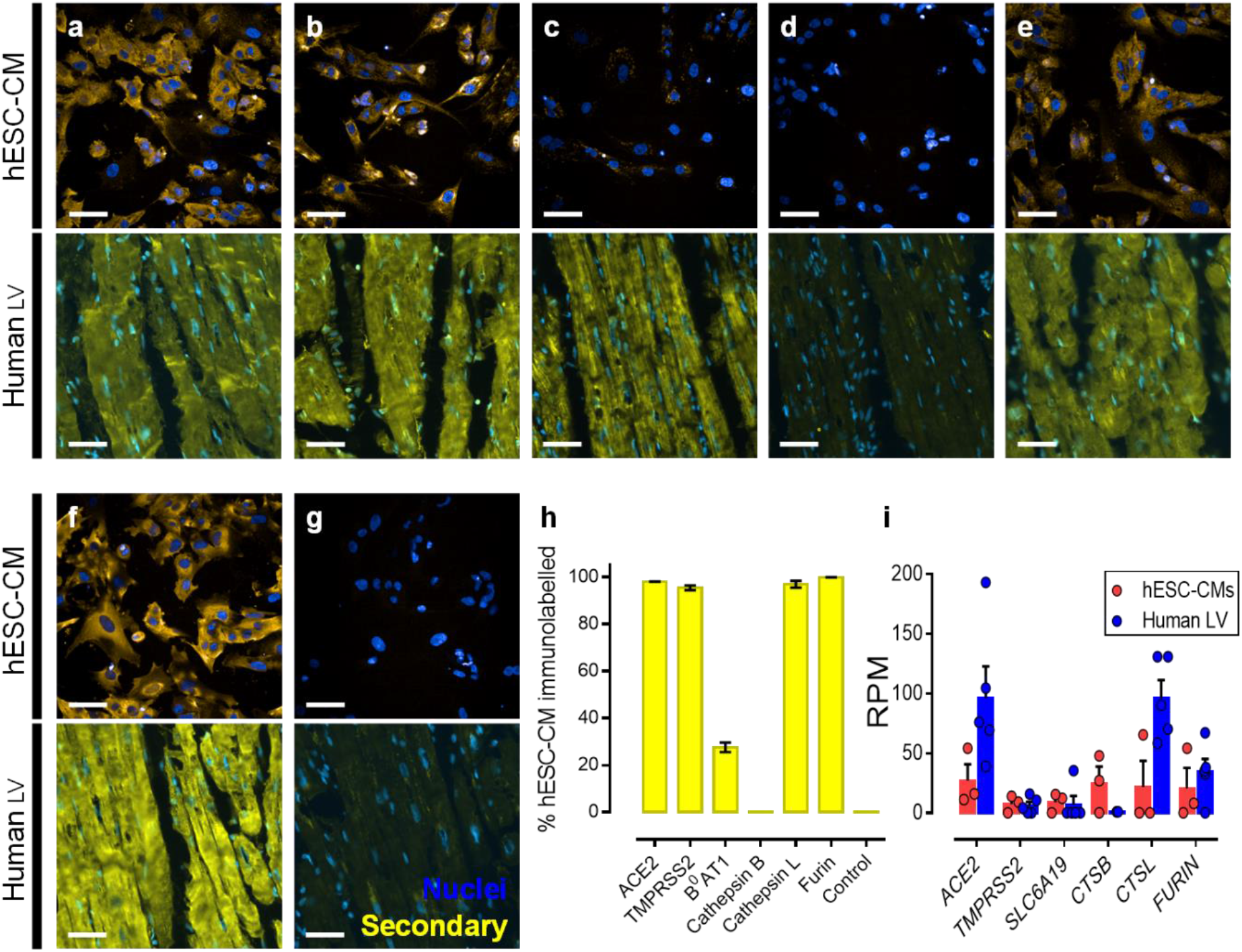
Detection of host cell proteins and genes associated with SARS-CoV-2 viral infection. **a-f**, Representative fluorescent confocal images (n=3 independent experiments performed in duplicate) of human embryonic stem cell-derived cardiomyocytes (hESC-CMs) (upper) and representative fluorescent images (n=6 from 6 different donors) of human left ventricle (Human LV) tissue sections (lower). Both cells and tissue were fixed with 4 % formaldehyde and immunolabelled with primary antibodies raised against ACE2 (**a**), TMPRSS2 (**b**), B^0^AT1 (**c**), cathepsin B (**d**), cathepsin L (**e**), and furin (**f**), before visualisation with secondary antibody conjugated to Alexa Fluor 555 (yellow) and Hoechst 33342 nuclear marker (blue). **g**, shows control cells (upper) and tissue (lower) treated with secondary antibody only and Hoechst 33342 nuclear marker. Scale bars show 50 μm. **h**, Graphical data showing the percentage of the observed hESC-CM population positively immunolabelled (above background) after visualisation with secondary antibody targeting primary antibodies raised against the outlined protein targets. **i**, Graphical data showing the reads per million (RPM) ± SEM for expression of viral entry and processing genes in hESC-CMs (n=7 replicates across 3 distinct differentiations) and human left ventricle (n=5 individuals). *SLC6A19, CTSB*, and *CTSL* are the genes that encode B^0^AT1, cathepsin B, and cathepsin L, respectively.

After demonstrating the presence of the protein complement required for SARS-CoV-2 viral entry in hESC-CMs, we infected these cells with SARS-CoV-2 and successfully showed titre- and time-dependent levels of infection (Supplementary Fig. 1).

Next, a drug screening platform was designed (Fig. 2a) using the beating hESC-CMs in conjunction with a SARS-CoV-2 spike pseudotyped GFP-expressing lentivirus to infect the cell model^9,18^. Infected differentiated cardiomyocytes were visualised in 96 well plates using a high content screening system (Opera Phenix; PerkinElmer), that allows for fast acquisition of fluorescent confocal images and subsequent quantification of viral entry into the hESC-CMs (Fig. 2b-e). Cells incubated with virus in media or DMSO (0.6 %) showed high rates of infection – 64.9 ± 4.2 % and 61.8 ± 5.8 % respectively of the observed cell population).

**Fig 2.**
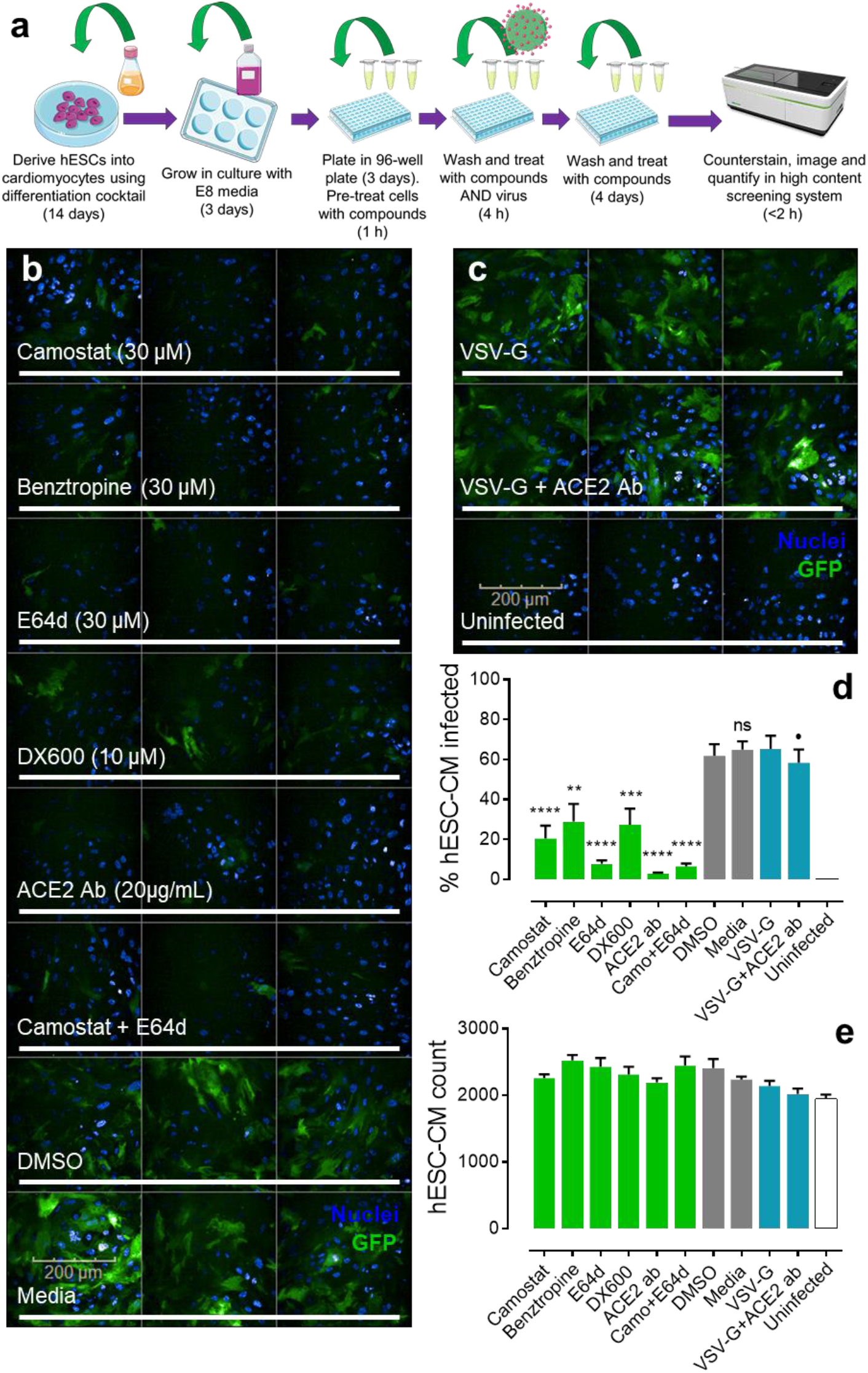
Pseudotyped lentiviral infection in hESC-CMs. **a**, Schematic showing the experimental workflow in brief for generating human embryonic stem cell-derived cardiomyocytes (hESC-CMs) and taking them into the pseudotyped lentiviral infection drug screen before conducting quantitative imaging (see methods for further details). Schematic was generated using templates from Servier Medical Art. **b**, Representative fluorescent confocal images (n=2 independent experiments performed in triplicate) of hESC-CMs pre-treated with small molecule inhibitors (camostat, benztropine, E64d), peptide antagonist (DX600), or antibody (ACE2 Ab) targeting protein components involved in SARS-CoV-2 infection. Control cells were treated with DMSO (0.6 %) or media. Cells were treated with drugs for 1 h before incubation with SARS-CoV-2 spike pseudotyped GFP-expressing (green) lentivirus for 4 h. After removal of viral particles, cells were washed and maintained in the presence of drugs for 5 days before fixation with 4 % formaldehyde and staining with Hoechst 33342 nuclear marker (blue). Scale bar shows 200 μm. **c**, Representative fluorescent confocal images (n=2 independent experiments performed in triplicate) of control human embryonic stem cell-derived cardiomyocytes (hESC-CMs) treated with VSV-G pseudotyped GFP-expressing (green) lentivirus, in the absence (upper) or presence (middle) of antibody (ACE2 Ab). Uninfected controls were not treated with viral particles (bottom). Again, cells were stained with Hoechst 33342 nuclear marker. **d**, Graphical data showing the percentage of observed hESC-CMs infected with either SARS-CoV-2 spike or VSV-G (control) pseudotyped lentivirus in the presence of drugs or DMSO (0.6 %) as indicated. Uninfected cells were not treated with viral particles. ** = p<0.005; *** = p<0.0005; **** = p<0.00005; and ns = no significant difference (as determined by one-way ANOVA) for each condition versus the DMSO treated control cells. · = no significant difference for condition versus the VSV-G control. **e**, Graphical data showing the overall count of observed hESC-CMs for each condition, as indicated. No condition showed a count significantly different (as determined by one-way ANOVA) from the DMSO treated control cells.

In wells treated with drugs targeting the proteins involved in SARS-CoV-2 viral entry and processing, we observed significant reductions in levels of infection. An ACE2 antibody, that has been shown previously to neutralize pseudotyped virus and SARS-CoV-2 infection^9^, was effective in our drug screen, significantly reducing infection level to 2.9 ± 0.4 %, as was DX600, the ACE2 peptide antagonist, which reduced infection to 20.5 ± 6.5 % in the observed cells. This highly selective peptide has not been tested previously as a viral entry inhibitor. DX600 forms multiple interactions with the catalytic site, that is distinct from the receptor binding domain of the virus, but may be sufficient to change the conformation of ACE2, or induce steric hindrance, to inhibit spike binding.

The ACE2 antibody did not significantly reduce the rate of infection with control vesicular stomatitis virus (VSV-G) pseudotyped lentivirus (Fig. 2b-d) (65.2 ± 6.7 % in the absence of the antibody and 58.3 ± 6.7 % in its presence), providing evidence for ACE2 dependency of the SARS-CoV-2 spike pseudotyped lentiviral entry. Uninfected cells were treated with media that did not contain pseudotyped lentivirus particles and showed little to no infection. These results confirm the requirement of expression of ACE2 in these cells for receptor recognition and viral entry, with no evidence for direct membrane fusion, as is common with other viruses.

Camostat and E64d, inhibitors of the accessory proteins TMPRSS2 and cathepsins respectively, significantly reduced infection levels to 20.5 ± 6.5 % and 7.8 ± 1.7 %. A mix of camostat and E64d effectively reduced infection rate as well (6.5 ± 1.5 %) but this was not significantly different to E64d treatment alone. These results suggest that inhibition of viral entry can be achieved at distinct steps of viral entry and processing. Importantly, camostat is already clinically approved in Japan where it is used to treat chronic pancreatitis, and is considered well tolerated and safe^19,20^. Additionally, camostat has been previously confirmed as an effective inhibitor of pseudotyped virus and SARS-CoV-2 infection^9^. Interestingly, benztropine, a small molecule inhibitor of the potential ancillary protein for viral entry, B^0^AT1, also successfully reduced pseudotyped virus infection levels (28.9 ± 8.8 %). The compound is used clinically as an adjunct in the therapy of all forms of parkinsonism but the precise mechanisms behind its inhibition of viral infection require further investigation. It is important to note that B^0^AT1 is found associated with ACE2 in high abundance in the gastrointestinal tract^13^.

Cell counts in the observed regions in the Opera Phenix (Fig. 2e), confirmed that hESC-CM number was not significantly altered in the presence of the compounds tested versus the media or DMSO (0.6 %) treated cells, confirming that treatments were unlikely to be toxic. Uninfected cells and those cells treated with pseudotyped lentivirus were also all present in comparable numbers, suggesting the viral inoculation was not inherently toxic. Cardiomyocytes were confirmed as beating visually, and a troponin-T cardiac specific marker demonstrated that hESC-CM populations were >95 % pure (Supplementary Fig. 2).

Our results have identified and validated a qualitative and quantitative screen, crucially using hESC-derived beating cardiomyocytes, where these clinically relevant cells are not rate limiting and widely available. We have demonstrated these are also viable and can be handled reliably in 96 well plates, the minimum format usually required for high throughput screens. hESC-CMs are shown here to allow for identification of current clinically approved medicines and discovery of novel compounds directed against the host protein targets used by SARS-CoV-2 to infect cells. The pseudotyped lentiviral system can also be handled at intermediate Biosafety levels, or containment levels of 2 meaning independent research groups will be able to use this platform without the need for higher-level facilities. In our studies, the methods described have allowed us to identify benztropine and DX600 as novel inhibitors of SARS-CoV-2 viral entry, alongside confirming the inhibitory effects of camostat, E64d, and ACE2 antibody. New medicines will be required for prophylactic treatment for those where vaccination against SARS-CoV-2 is contraindicated, and for the need to protect the heart against acute damage during hospitalization, as well as during the prolonged post-viral recovery observed in certain individuals. Furthermore, vaccine selection pressure is likely to result in resistant strains and a screening campaign is needed to add inhibitors of viral entry to our armamentarium, a strategy that has proved remarkably successful in other instances such as maraviroc treatment in HIV cases. Our results provide a platform to generate high quality data in pre-clinical studies to justify translational research in animal models of acute respiratory distress syndrome and the repurposing of current drugs for clinical trials.

References limited to twenty to comply with Instructions to Authors. For comprehensive reviews on SARS-CoV-2 host protein drug targets, see references 13 and 14. For human cardiovascular stem cell derivatives and their uses in experimental medical research, see references 16 and 17.

## Acknowledgments

E.J.D.G. SARS-CoV2/human/Liverpool/REMRQ0001/2020 was a kind gift from Lance Turtle (University of Liverpool) and David Matthews and Andrew Davidson (University of Bristol).

## Author contribution

T.L.W. and M.T.C. designed and carried out experiments and data analysis; R.G.C.M. carried out experiments, performed RNAseq experiments and data analysis; E.L.R. performed RNAseq experiments and analysis; S.B. and J.R.Z. designed and generated pseudotyped lentiviral constructs; E.J.D.G. performed SARS-CoV-2 viral infections, imaging and analysis; G.S. contributed imaging expertise and facilities; R.E.K. and D.N. provided laboratory support; P.J.L. and S.S. supervised experiments and contributed grant support and facilities; J.J.M. and A.P.D. designed and supervised experiments, performed data analysis, contributed grant support and facilities. All authors contributed to the writing and/or review of the manuscript.

## Competing Interests

The authors declare no competing interests.

## Methods

### Short Methods

#### hESC Culture and Differentiation to Cardiomyocytes

Pluripotent H9 hESCs (WiCell) were maintained in culture as described previously (Cheung *et al*., 2014)^21^. hESCs were grown as colonies in Essential 8 medium (E8), supplemented with fibroblast growth factor 2 (FGF2,25 ng/ml, Qkine Ltd) and TGF-b (1.74 ng/ml, R&D Biotechne), on culture plates previously coated with vitronectin (Stem cell technologies). A previously optimised protocol was used to induce differentiation of hESCs to spontaneously contracting cardiomyocytes (adapted from Mendjan *et al*., 2014)^22^. Briefly, differentiation was directed towards mesoderm lineage by culturing in CDM-BSA media supplemented with FGF2 (20 ng/ml), Activin-A (50 ng/ml) and Bone Morphogenetic Protein 4 (BMP4, 10 ng/ml, Bio-Techne) plus phosphoinositide 3-kinase inhibitor Ly294002 (10 µM, Promega). After 42 h incubation at 37°C, media was replaced with CDM-BSA supplemented with FGF2 (8 ng/ml), BMP4 (10 ng/ml), retinoic acid (1 µM, Sigma Aldrich) and the WNT signalling pathway inhibitor IWR1-endo (1 ng/ml, R&D Systems), which was refreshed after 48 h. After a further 48 h, media was changed for CDM-BSA supplemented with FGF2 (8 ng/ml) and BMP4 (10 ng/ml). Media was replaced with normal CDM-BSA after 48 h, and cardiomyocytes were maintained in CDM-BSA (refreshed every other day) until robust beating was established. Differentiation efficiency using this protocol has previously been established as around 85-90% cardiomyocytes, as evidenced by troponin positive percentage by flow cytometry (as previously described in Bargehr *et al*., 2019)^23^. If the percentage of Troponin positive cells was lower than 85%, cells were subjected to lactate selection as previously described. Briefly, cells were cultured in glucose free media (DMEM) for 4 days in the presence of Sodium-Lactate (SIGMA). At the end of the selection purity was then re-evaluated by flow cytometry. Independent replicates were considered to be cardiomyocytes generated from distinct differentiations.

### Flow Cytometry

hESC-derived cardiomyocytes were collected as pellets and resuspended in Fixation/Solubilization solution (BD Cytofix/Cytoperm Fixation/Permeabilization Kit, Biosciences) for 20 mins at 4°C. Cells were then pelleted by centrifugation, washed three times in phosphate buffered saline containing 0.1% BSA and 2 mM EDTA (PBE), then resuspended in PBE containing FITC-conjugated antibody specific for cardiac troponin T (Miltenyi Biotec, cat no. 130-119-575) at a concentration of 1:50 and incubated for 2 h at 4°C. Three washes were performed using PBE and then cells were resuspended in PBE and run on LSRFortessa X-20 Flow Cytometer (BD Biosciences). Analysis was performed using FlowJo software (BD Biosciences). This research was supported by the Cambridge NIHR BRC Cell Phenotyping Hub.

### SARS-CoV-2 virus stock

The SARS-CoV-2 virus used in this study is the clinical isolate named “SARSCoV2/human/Liverpool/REMRQ0001/2020”^24,25^. In total, the stock used was passaged three times in VeroE6 cells, once in Caco2 cells, and once in Calu3 cells. The titre of the virus stock was determined by TCID_50_ (Median Tissue Culture Infectious Dose) assay on Huh7 cells transduced with an ACE2 expression vector.

### SARS-CoV-2 immunostaining

SARS-CoV-2 infected cells were fixed in 2% formaldehyde for 30 mins and permeabilized with BD Perm/Wash buffer (BD Biosciences, 554723). The primary antibody used was sheep anti-SARS-CoV-2 nucleocapsid antibody (DA114, MRC-PPU), which was visualised with AF488 conjugated donkey anti-sheep antibody (Jackson ImmunoResearch #713-545-147)

### Pseudotyped virus production

#### Cell lines

HEK293T cells (Human embryonic kidney, Lehner laboratory stocks) were cultured in Iscove’s Modified Dulbecco’s Media (Sigma) supplemented with 1% GlutaMAX, 10% FCS, 100 units/ml penicillin and 100 µg/ml streptomycin (all from Thermo Fisher), at 37°C in 5% CO2. Cells were confirmed mycoplasma negative (MycoAlert, Lonza).

#### Plasmids

Plasmid pCG1-SARS-CoV-2 Δ19 D614G HA, encoding the SARS-CoV-2 spike protein carrying a D614G change, with the C-terminal 19 amino acids replaced with an HA Tag, was generated from pCG1-SARS-2-S (a kind gift from M. Hoffmann^9^ by PCR (Phusion polymerase (NEB)) using primer pairs; (1) CoV2optFor (TTGTATCGGATCCACCATGTT CGTGTTTCTGGTGCTGCTG) with CoV2D614Gr (CAGTTCACGcCCTGGTACAGCAC TGCCAC) and (2) CoV2D614Gf (GTACCAGGgCGTGAACTGTACCGAAGTGCC) with CoV2optD19HArev (ATCCCGATCTAGATCAAGCGTAATCTGGAACATCGTATGGGT AACTTCCGCAGCAGCTGCCACAGCTACA) to generate overlapping amplification products, that were gel purified and combined as template in a second PCR reaction using primer pair; CoV2optFor and CoV2optD19HArev. The final amplification product was digested and re-cloned into the pCG1 vector using BamHI-XbaI sites. The HIV-1 based GFP-expressing lentiviral vector pHRSIN CSGW eGFP was a kind gift from Y. Ikeda^26^.

Concentrated pseudotyped lentivirus production

Pseudotyped lentiviral supernatants were generated following co-transfection of HEK293T cells with pHRSIN CSGW eGFP plus the packaging plasmid pCMVΔR8.91 and either pMD.G (VSV-G) or pCG1-SARS-CoV-2 Δ19 D614G HA spike plasmid, using TransIT-293 transfection reagent (Mirus) according to the manufacturer’s recommendations. Viral supernatants were harvested 48 h post transfection and cell debris was removed with a 0.45-μm syringe filter, prior to concentration (approximately 10-fold) using a Vivaspin 20 MWCO 100kDa centrifugal concentrator (Sartorius) according to manufacturer’s recommendations.

### Pseudotyped virus infection of hESC-derived cardiomyocytes

For infection of hESC-derived cardiomyocytes, cells in CellCarrier-96 Ultra Plates (PerkinElmer) were incubated for 4 h with pseudotyped viral stock at the desired MOI in media. In drug screens, cells were pre-treated for 1 h before infection with either camostat, benztropine, or E64d at a final concentration of 30 μM in media; DX600 at a final concentration of 10 μM in media; ACE2 antibody (AF933; R&D Systems) at 20 μg/mL; a mix of camostat + E64d at a final concentration of 30 μM each; or DMSO at 0.6% (equivalent of the highest concentration included in drug dilutions). Initial toxicity tests demonstrated that these concentrations were not toxic after 5 days. Cells were then washed 3x with PBS to remove infectious lentivirus particles, before replenishment with the media outlined above, maintained in the presence of the respective drug treatments for 4-5 days. Viral infection was monitored using an EVOS Cell Imaging System to look for green fluorescence protein (GFP) as a marker of viral presence in cells. See below for details on visualisation of these cells in the Opera Phenix High Content Screening System.

### Opera Phenix High Content Screening System (PerkinElmer) Methods

For immunocytochemistry, hESC-derived cardiomyocytes in CellCarrier-96 Ultra Plates (PerkinElmer) were washed 3x with 100 μL PBS before fixation with 50 μL buffered (pH 6.9) 4% formaldehyde solution for 20 mins. Subsequently, non-specific staining was blocked with PBS containing 10% donkey sera for 2 h at room temperature. Cells were aspirated and incubated overnight at 4°C with either: primary goat polyclonal antibody to Human ACE-2 (AF933; R&D Systems; 1:100); primary rabbit monoclonal antibody to TMPRSS2 (ab92323; Abcam; 1:500); primary rabbit monoclonal antibody to B^0^AT1 (ab180516; Abcam; 1:300); primary rabbit monoclonal antibody to cathepsin B (ab125067; Abcam; 1:100); primary rabbit polyclonal antibody to cathepsin L (ab203028; Abcam; 1:100); or primary rabbit polyclonal antibody to furin (ab3467; Abcam; 1:500), all prepared in PBS with 1% donkey sera, 0.1% Tween-20, and 3.3 mg/mL bovine serum albumin. Buffer treated controls were incubated with PBS with 1% donkey sera, 0.1% Tween-20, and 3.3 mg/mL bovine serum albumin alone. After 24 h, cells were washed 3x with 100 μL PBS with 0.1% Tween-20 before incubation with the secondary polyclonal Donkey Anti-Goat IgG H&L antibody conjugated to Alexa Fluor 555 (ab150130; Abcam; 1:200) or Donkey Anti-Rabbit IgG H&L antibody conjugated to Alexa Fluor 555 (ab150066; Abcam; 1:200) prepared at 0.01 mg/mL in PBS with 1% donkey sera, 0.1% Tween-20, and 3.3 mg/mL bovine serum albumin, for 1 h at room temperature. Cells were washed 3x with 100 μL PBS before incubation with Hoechst 33342 nuclear stain (H3570; Invitrogen) prepared at 10 μg/mL in PBS for 15 mins at room temperature in the dark. After a final 3x washes with 100 μL PBS, cells were maintained in 100 μL PBS for imaging.

For viral infection experiments, pseudotyped virus treated hESC-derived cardiomyocytes in CellCarrier-96 Ultra Plates (PerkinElmer) were washed 3x with 100 μL PBS before complete submersion in 2-4% formaldehyde solution for 30 mins. After fixation, hESC-derived cardiomyocytes plated in CellCarrier-96 Ultra Plates (PerkinElmer) were washed 3x with 100 μL PBS before incubation with Hoechst 33342 nuclear stain (H3570; Invitrogen) prepared at 10 μg/mL in PBS for 15 mins at room temperature in the dark. Cells were washed a further 3x with 100 μL PBS and maintained in 100 μL PBS for imaging.

Fluorescent confocal images of cells were acquired using the Opera Phenix High Content Screening System (PerkinElmer) microscope with a 40x/NA1.1 water immersion objective. The Opera Phenix is a dual camera, sequential, multiple laser, spinning disk confocal microscope that acquires images in a fast and gentle manner to minimise fluorescent bleaching. For immunocytochemistry, excitation/emission laser and filter sets for two fluorescent channels were used: 405/435-480 nm (blue) for the Hoechst 33342 nuclear stain, and 561/570-630 nm (yellow) for the Donkey Anti-Goat IgG H&L antibody conjugated to Alexa Fluor 555 (ab150130; Abcam) or Donkey Anti-Rabbit IgG H&L antibody conjugated to Alexa Fluor 555 (ab150066; Abcam). For viral infection experiments, excitation/emission laser and filter sets for two fluorescent channels were used: 405/435-480 nm (blue) for the Hoechst 33342 nuclear stain, and 488/500-550 nm (green) for GFP or the donkey anti-sheep antibody conjugated to Alexa Fluor 488 (Jackson ImmunoResearch #713-545-147). Laser intensities, for blue (405 nm) and green (488 nm) channels respectively, were both set to 50% transmission, with a 50 ms exposure time. Opera Phenix hardware focus ensures a consistent focal depth across the CellCarrier-96 Ultra Plate is achieved. Scale bars show 50 μm unless specified otherwise.

In-built Opera Phenix quantitation was performed using Harmony High-Content Imaging and Analysis Software (PerkinElmer). For analysis, images were filtered, and the contrast of the Hoechst nuclear blue channel was increased 5-fold to ensure cells were distinguished from background and to assist in finding the outer limits of individual cells. Fluorescence intensities were then calculated for every observed cell in the selected population and were deemed positive where whole cell fluorescence (averaged over individual cell area) was ≥130 fluorescent units, where background was determined as ∼113 fluorescent units.

### Slide Scanner Axio Scan.Z1 (Zeiss) Methods

Surgical samples of human tissue were obtained with informed consent from Royal Papworth Hospital Research Tissue Bank and ethical approval (05/Q104/142). Tissues were snap frozen in liquid nitrogen before storage at −80°C. Tissue samples from humans (n = 6 individuals) were cut, using a cryostat (−20°C to −30°C), into 10 μm sections and thaw mounted onto slides before return to storage at −80°C. On the day of experimentation, frozen tissue sections were thawed for 20 mins at room temperature (21°C), encircled with a hydrophobic pen, and subsequently rehydrated with PBS. After 3x washes with PBS, tissue sections were fixed with buffered (*p*H 6.9) 4% formaldehyde solution for 20 mins. Following a further 3x washes with PBS, non-specific binding was blocked with PBS containing 10% donkey sera for 2 h at room temperature. Sections were then incubated overnight at 4°C with either: primary goat polyclonal antibody to Human ACE-2 (AF933; R&D; 1:100); primary rabbit monoclonal antibody to TMPRSS2 (ab92323; Abcam; 1:500); primary rabbit monoclonal antibody to B^0^AT1 (ab180516; Abcam; 1:300); primary rabbit monoclonal antibody to cathepsin B (ab125067; Abcam; 1:100); primary rabbit polyclonal antibody to cathepsin L (ab203028; Abcam; 1:100); or primary rabbit polyclonal antibody to furin (ab3467; Abcam; 1:500), all prepared in PBS with 1% donkey sera, 0.1% Tween-20, and 3.3 mg/mL bovine serum albumin. After 24 h, sections were washed 3x with PBS with 0.1% Tween-20 before incubation with the secondary polyclonal Donkey Anti-Goat IgG H&L antibody conjugated to Alexa Fluor 555 (ab150130; Abcam; 1:200) or Donkey Anti-Rabbit IgG H&L antibody conjugated to Alexa Fluor 555 (ab150066; Abcam; 1:200) prepared at 0.01 mg/mL in PBS with 1% donkey sera, 0.1% Tween-20, and 3.3 mg/mL bovine serum albumin, for 1 h at room temperature. Tissue sections were washed again 3x with PBS before incubation with Hoechst 33342 nuclear stain (H3570; Invitrogen) prepared at 10 μg/mL in PBS for 15 mins at room temperature in the dark. Following a further 3x washes with PBS, slides were incubated for 1 h at room temperature in the dark with a cardiac troponin-T antibody conjugated to APC (130-120-543; Miltenyi Biotec) prepared at a 1:100 dilution in PBS. After a final 3x washes with PBS, slides were blotted dry with lint-free tissue, mounted with ProLong Gold Antifade Mountant (P36930; Invitrogen), covered with a cover slip, and left overnight at room temperature in the dark to dry.

Automated fluorescent images (16 bit, 0.325 x 0.325 μm scaling per pixel) were acquired using a Slide Scanner Axio Scan.Z1 (Zeiss) microscope with a Plan-Apochromat 20x/NA0.8 M27 objective lens connected to a Hamamatsu Orca Flash camera. The system uses an LED light source, which provides more consistent intensity over time. After using live bright field imaging to find tissue on the slides, three fluorescent channels were used, with low exposure times to minimise bleaching of the sample. The first (blue channel) used an LED-Module 385 nm light source set at 10% intensity and 10 ms exposure time at a depth of focus of 1.45 μm to illuminate the Hoechst nuclear stain (max excitation and emission of 361 and 497 nm respectively). The second (yellow channel) used an LED-Module 567 nm light source set at 80% intensity and 30 ms exposure time at a depth of focus of 1.88 μm to illuminate the Donkey Anti-Goat IgG antibody or Donkey Anti-Rabbit IgG H&L antibody, both conjugated to Alexa Fluor 555 (max excitation and emission of 555 and 580 respectively for both). The third (far red channel) used an LED-Module 630 nm light source set at 50% intensity and 20 ms exposure time at a depth of focus of 2.09 μm to illuminate the cardiac troponin-T antibody conjugated to APC (max excitation and emission of 650 and 660 respectively). Filters for wavelengths of 430-470, 600-660, and 670-710 nm were used for the blue, yellow, and red channels, respectively. Acquired images were saved and visualised using ZEN software (Zeiss). Scale bars show 50 μm unless specified otherwise.

### Statistical Analyses

All quantitative data are expressed as mean ± SEM. The *n* values are stated in the methods and/or the figure legends. Statistical one-way ANOVA tests with Dunnett’s correction for multiple comparisons were used. A *p* value of <0.05 was determined as significant. Graphical presentation and statistical analyses were performed using GraphPad Prism version 6.07 for Windows (GraphPad Software, La Jolla, California, USA).

### RNA Extraction for RNA-Seq

Total RNA extraction was performed using TRIzol reagent (Invitrogen) according to the manufacturer’s protocol. Briefly, for iPSC-derived cardiomyocytes, cell pellets were thawed and 1 mL TRIzol reagent and incubated for 5 mins to allow nucleoprotein dissociation. For human heart, tissue was homogenized mechanically in 1 mL TRIzol reagent using a Polytron and incubated for 5 mins to allow nucleoprotein dissociation. 180 µl of chloroform was added to each sample, mixed thoroughly and incubated for 3 mins at room temperature. Samples were then centrifuged at 12,000 xg for 15 mins at 4°C to promote phase separation. The RNA-containing upper aqueous phase was transferred to a fresh tube. Isopropanol (500 µl) was added and incubated for 10 mins to precipitate RNA. Samples were then centrifuged at 10,000 xg for 10 mins at 4°C, supernatant discarded, and RNA precipitate collected as a pellet. Pellets were resuspended in 1 mL 75 % ethanol, vortexed briefly and spun at 7,500 xg for 5 mins at 4°C to wash the RNA. The supernatant was discarded, and pellets allowed to air dry for 10 mins at room temperature before being resuspended in 20 µl RNase-free water. RNA concentration was determined using a NanoDrop 1000 (ThermoFisher), and RNA samples subsequently stored at −70°C prior to RNA sequencing library preparation.

### RNA Processing and Sequencing

Quality control: RNA quality was verified using the TapeStation RNA ScreenTape (Agilent). All control HLV and stem cell RNA samples had RINe 7.14 – 9.0 (7.8 +/-0.3). Q.C. was performed at the Cambridge Genomics Services (Department of Pathology, University of Cambridge). Ribosomal RNA was removed using NEBNext® rRNA Depletion Kit (Human/Mouse/Rat) (New England Biolabs) according to the manufacturer’s instructions with 6 µl total RNA used as input per sample. Total stranded RNA-sequencing library preparation and quality control: Total stranded RNA-sequencing libraries were generated using the CORALL Total RNA-Seq Library Prep Kit (Lexogen) according to manufacturer’s instructions, with 15 PCR cycles used for the final amplification step and passed through quality control using a 2100 Bioanalyzer (Agilent). Both quality control and sequencing was carried out at Babraham Institute Next Generation Sequencing facility. 15 RNA-seq libraries were sequenced per lane of a HiSeq2500 (Illumina) as 100bp Single-End sequencing runs.

### Data Analysis

Processing and analysis of next generation sequencing data: All next generation sequencing data were aligned using HiSAT2 (http://daehwankimlab.github.io/hisat2/) to the Homo sapiens reference genome GRCh38/hg38. Reads were trimmed prior to alignment using Trim Galore, using Phred quality score for base calling cut-off of 20, corresponding to a maximum error of 1 in 100 bases and with a maximum trimming error rate of 0.1 (http://www.bioinformatics.babraham.ac.uk/projects/trim_galore/)^27^. Trimmed and aligned sequence files were imported as BAM files into SeqMonk (v1.42.0) for visualization and analysis (http://www.bioinformatics.babraham.ac.uk/projects/seqmonk/).

RNA-seq analysis for differential gene expression: Sequencing reads were quantified by read count quantitation and global normalisation performed to total read count for each replicate and expressed as reads per million (RPM). A screen of a number of different cell types from different species found inconsistency between samples using RPKM and that using RPKM for normalisation did not respect the invariance property^28^. Differential gene expression analysis was performed using the R-based software DESeq2^29^. Raw (non-log transformed) read counts were used as input and global normalization performed to total library size. RPM values ≥ 1.0 were deemed to be above noise. Benjamini-Hochberg correction for multiple testing was used with false discovery rate of 5 %.

## Notes

### Competing Interest Statement

The authors have declared no competing interest.

