## Supplementary Figure 1 for "Human embryonic stem cell-derived cardiomyocytes express SARS-CoV-2 host entry proteins: screen to identify inhibitors of infection"

### Slide 1
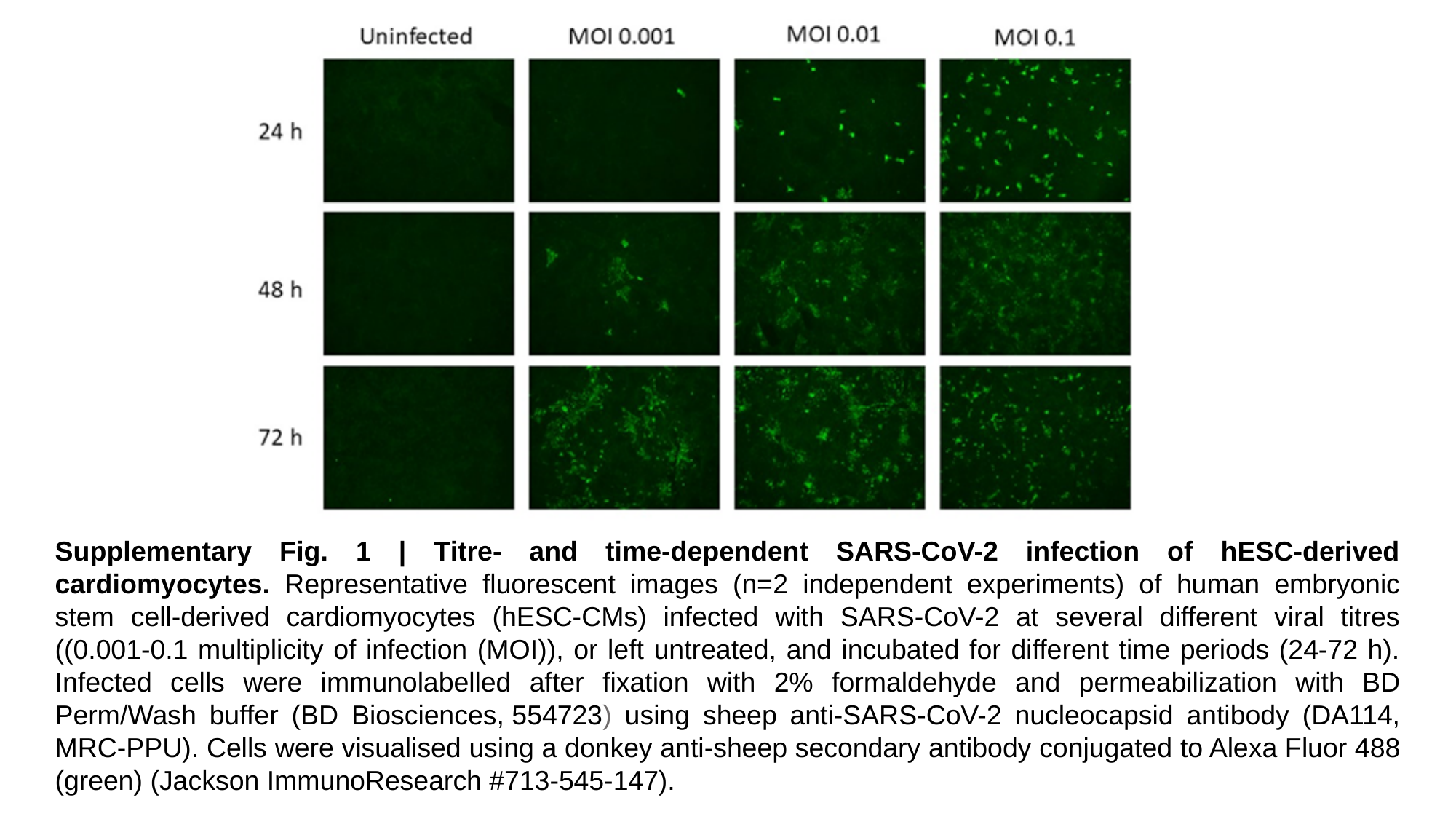

Supplementary Fig. 1 | Titre- and time-dependent SARS-CoV-2 infection of hESC-derived cardiomyocytes. Representative fluorescent images (n=2 independent experiments) of human embryonic stem cell-derived cardiomyocytes (hESC-CMs) infected with SARS-CoV-2 at several different viral titres ((0.001-0.1 multiplicity of infection (MOI)), or left untreated, and incubated for different time periods (24-72 h). Infected cells were immunolabelled after fixation with 2% formaldehyde and permeabilization with BD Perm/Wash buffer (BD Biosciences, 554723) using sheep anti-SARS-CoV-2 nucleocapsid antibody (DA114, MRC-PPU). Cells were visualised using a donkey anti-sheep secondary antibody conjugated to Alexa Fluor 488 (green) (Jackson ImmunoResearch #713-545-147).
