## Supplementary Figure 2 for "Human embryonic stem cell-derived cardiomyocytes express SARS-CoV-2 host entry proteins: screen to identify inhibitors of infection"

### Slide 1
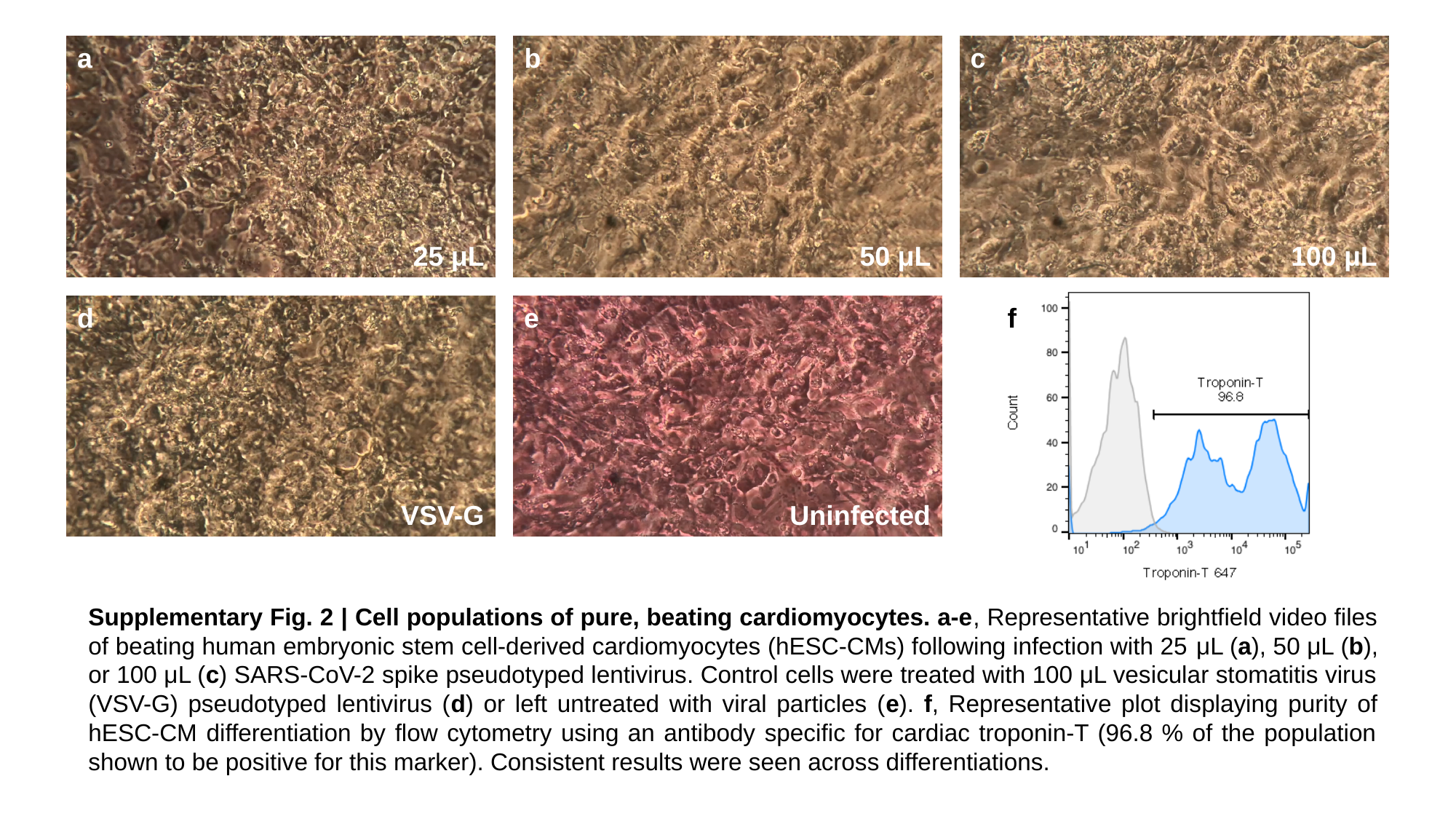

a
b
c
25 μL
50 μL
100 μL
d
e
f
VSV-G
Uninfected
Supplementary Fig. 2 | Cell populations of pure, beating cardiomyocytes. a-e, Representative brightfield video files of beating human embryonic stem cell-derived cardiomyocytes (hESC-CMs) following infection with 25 μL (a), 50 μL (b), or 100 μL (c) SARS-CoV-2 spike pseudotyped lentivirus. Control cells were treated with 100 μL vesicular stomatitis virus (VSV-G) pseudotyped lentivirus (d) or left untreated with viral particles (e). f, Representative plot displaying purity of hESC-CM differentiation by flow cytometry using an antibody specific for cardiac troponin-T (96.8 % of the population shown to be positive for this marker). Consistent results were seen across differentiations.
