## Supplementary Table 1 for "Human embryonic stem cell-derived cardiomyocytes express SARS-CoV-2 host entry proteins: screen to identify inhibitors of infection"

### Slide 1
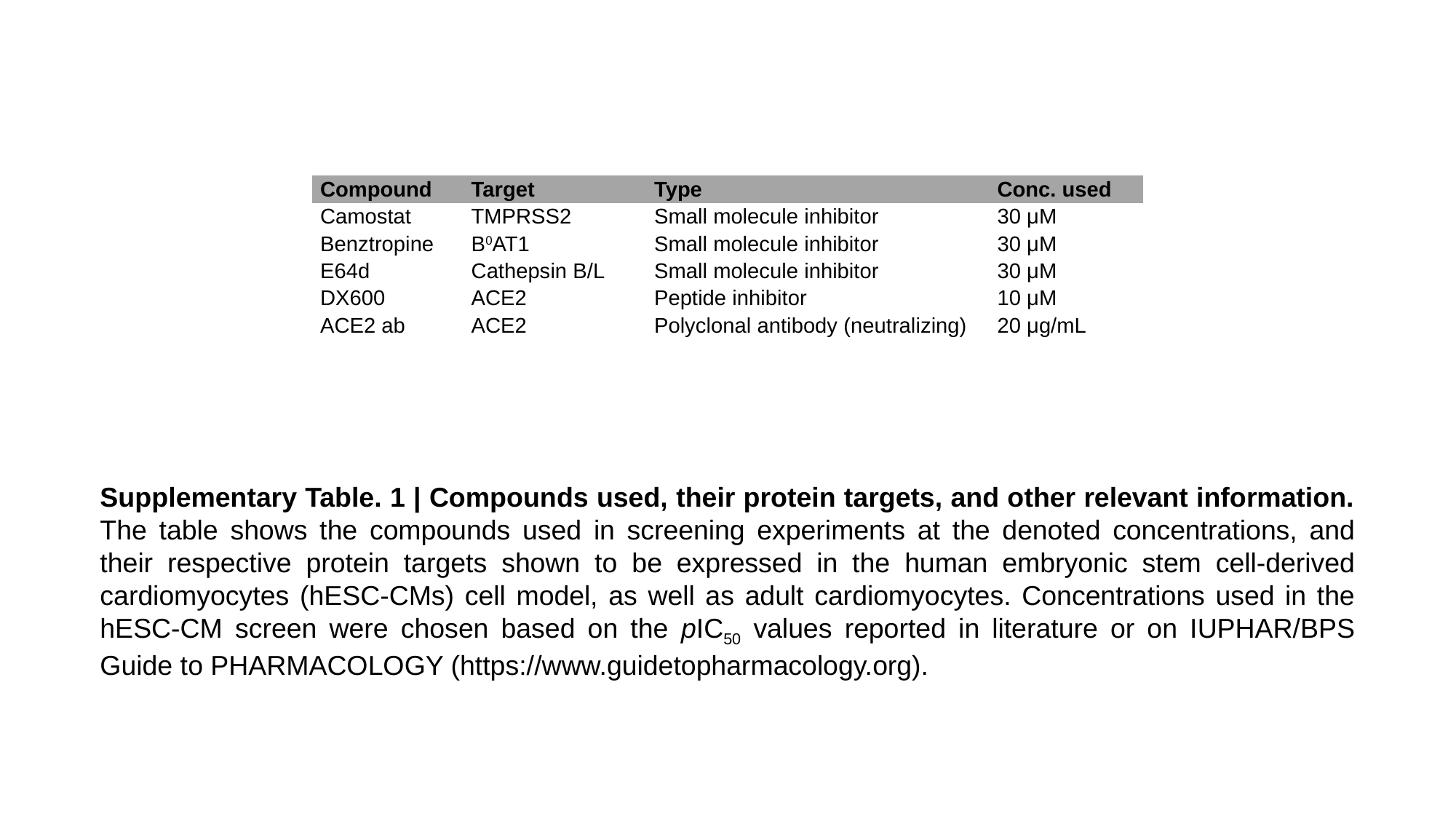

| Compound | Target | Type | Conc. used |
| --- | --- | --- | --- |
| Camostat | TMPRSS2 | Small molecule inhibitor | 30 μM |
| Benztropine | B^0^AT1 | Small molecule inhibitor | 30 μM |
| E64d | Cathepsin B/L | Small molecule inhibitor | 30 μM |
| DX600 | ACE2 | Peptide inhibitor | 10 μM |
| ACE2 ab | ACE2 | Polyclonal antibody (neutralizing) | 20 μg/mL |
Supplementary Table. 1 | Compounds used, their protein targets, and other relevant information. The table shows the compounds used in screening experiments at the denoted concentrations, and their respective protein targets shown to be expressed in the human embryonic stem cell-derived cardiomyocytes (hESC-CMs) cell model, as well as adult cardiomyocytes. Concentrations used in the hESC-CM screen were chosen based on the pIC_50_ values reported in literature or on IUPHAR/BPS Guide to PHARMACOLOGY (https://www.guidetopharmacology.org).
